# AI on animals: AI-assisted animal-borne logger never misses the moments that biologists want

**DOI:** 10.1101/630053

**Authors:** Joseph M. Korpela, Hirokazu Suzuki, Sakiko Matsumoto, Yuichi Mizutani, Masaki Samejima, Takuya Maekawa, Junichi Nakai, Ken Yoda

**Affiliations:** Graduate School of Information Science and Technology, Osaka University, Suita, Osaka 565-0871, Japan; Graduate School of Environmental Studies, Nagoya University, Nagoya, Aichi 464-8601, Japan; Graduate School of Dentistry, Tohoku University, Sendai, Miyagi 980-8575, Japan

## Abstract

Animal-borne data loggers, i.e., biologgers, allow researchers to record a variety of sensor data from animals in their natural environments (Hussey et al. 2015; Kays et al. 2015). This data allows biologists to observe many aspects of the animals’ lives, including their behavior, physiology, social interactions, and external environment. However, the need to limit the size of these devices to a small fraction of the animal’s size imposes strict limits on the devices’ hardware and battery capacities (Kays et al. 2015). Here we show how AI can be leveraged on board these devices to intelligently control their activation of costly sensors, e.g., video cameras, allowing them to make the most of their limited resources during long deployment periods. Our method goes beyond previous works that have proposed controlling such costly sensors using simple threshold-based triggers, e.g., depth-based (Watanuki et al. 2007; Volpov et al. 2015) and acceleration-based (Nishiumi et al. 2018; Brown et al. 2012) triggers. Using AI-assisted biologgers, biologists can focus their data collection on specific complex target behaviors such as foraging activities, allowing them to automatically record video that captures only the moments they want to see. By doing so, the biologger can reserve its battery power for recording only those target activities. We anticipate our work will provide motivation for more widespread adoption of AI techniques on biologgers, both for intelligent sensor control and intelligent onboard data processing. Such techniques can not only be used to control what is collected by such devices, but also what is transmitted off the devices, such as is done by satellite relay tags (Cox et al. 2018).

## INTRODUCTION

‘Bio-logging,’ i.e. the use of animal-borne sensors has revolutionized the study of animal behavior in the natural environments (Hussey et al. 2015; Kays et al. 2015). Although there have been extraordinary improvements in sensors and memories of the devices since the first logger was attached to a Weddell seal (Kooyman 1965), behavioral time-series data has been obtained with a simple strategy: continuous recording regardless of the researchers’ goals. For example, video loggers continue to shoot animal behavior and the surrounding environment including unimportant scenes, which consumes a large amount of power. Because the size of an animal-borne device is limited by the animal’s carrying capacity, ‘intelligent’ technology is needed for increasing the potential to apply bio-logging in a variety of research fields.

In this study, we propose the concept of AI-assisted biologgers that use low-cost sensors to automatically detect activities of interest, allowing them to conditionally activate high-cost sensors to target those activities. Although simple threshold-based camera trigger mechanisms are available, e.g., acceleration-based GPS triggers (Brown et al. 2012), it is difficult for biologists to capture complex activities of interest with these mechanisms due to the difficulty in creating rules for detecting complex activities using only simple thresholding.

These costs can vary depending on the application, with examples including the use of low cost (low bitrate) GPS data to control a high cost (high bitrate) microphone that normally would quickly fill the device’s storage, or the use of a low cost (low energy) acceleration sensor to control the use of a high-cost (energy consuming) camera. In this study, we focus on the second of these examples, using acceleration and GPS data to control the use of our logger’s energy consuming camera because biologists’ demand for animal-borne video cameras keeps increasing for decades (e.g., Rutz et al. 2007; Moll et al. 2007; Gómez-Laich et al. 2015).

Fig. 1 (a) shows an example of how such a logger can be used for seabirds, with the logger attached to the back of a seabird which is then released to roam freely in its natural environment. Fig. 1 (b) shows how this logger can continuously run its low-energy sensors (e.g., an accelerometer) and use these sensors’ data to detect important activities, such as foraging. Upon detecting such important activities, the logger can then activate its energy consuming sensor (i.e., a camera) to record the important activity. By doing so, the logger can limit its use of the energy-consuming sensor to times when it is most likely to capture the target activity, increasing its chances for success by extending the runtime of the logger. This contrasts with normal loggers that continuously run the energy-consuming sensors, causing them to quickly exhaust their batteries which limits their chances for successfully recording the target activities.

**Figure 1.**
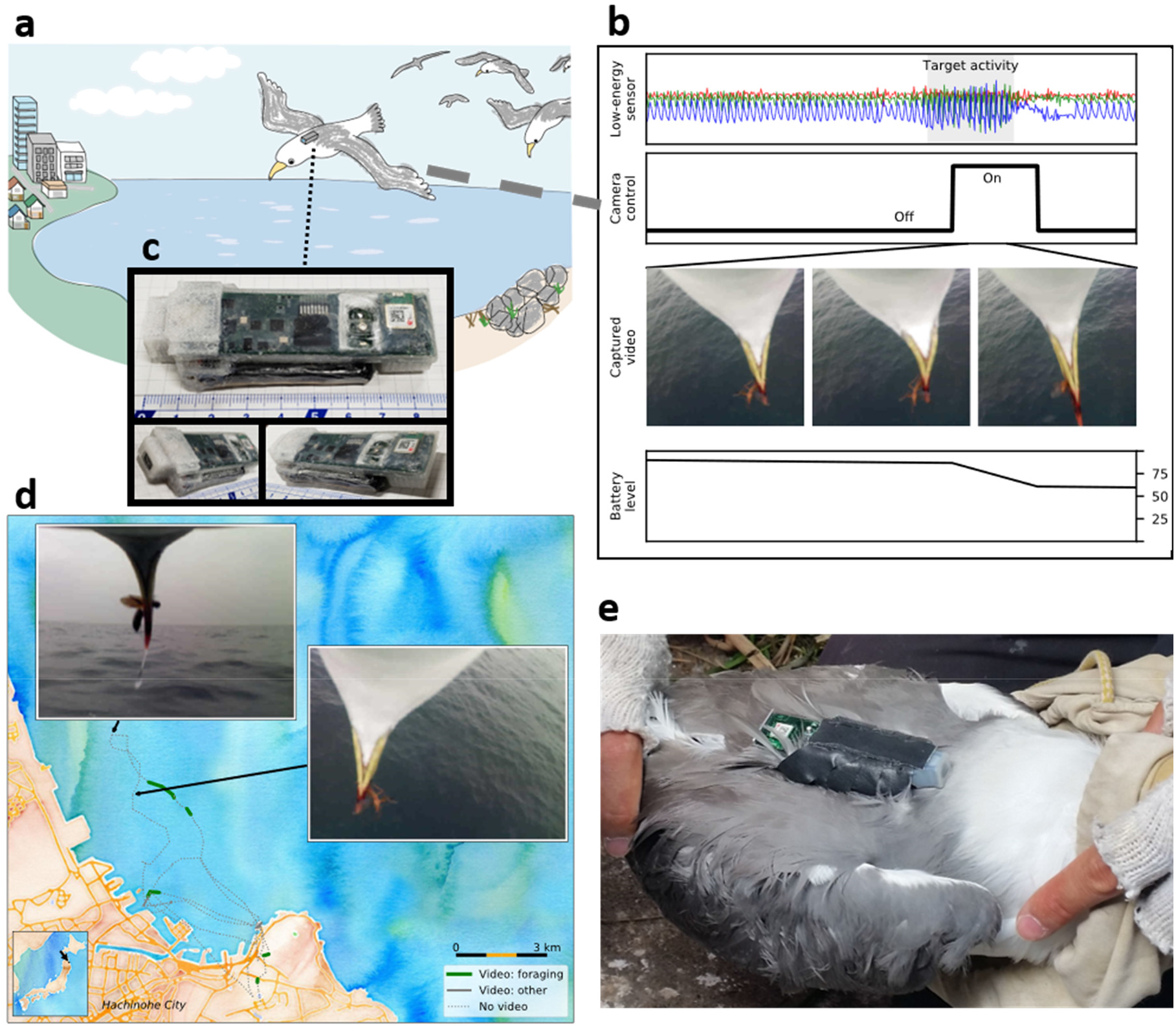
Biologging device used in this study. (a) Example deployment of biologger on a seabird in its natural environment. (b) Use of low-cost accelerometer to detect foraging activity and activate high-cost video camera for targeted collection. (c) Biologging device pictured with camera pointing to left, coated in waterproofing material for use on black-tailed gulls. Device measures 85 mm length x 35 mm width x 15 mm height and weighs approximately 27 g. (d) Example data collected by the biologging device from a single black-tailed gull from a colony near Hachinohe City, Japan. Green highlighted portions of GPS track indicate successful video recording of foraging behavior with inset images showing examples of insect predation captured by the device. (e) Attachment of biologging device in the field to the back of a black-tailed gull using Tesa tape.

In order to robustly detect animal activities using sensor data in the wild, we employ supervised learning to conduct activity recognition on board the logging devices. That is, we start by having a biologist label sensor data from low-energy sensors to identify the activities that he/she wants to record in advance. We then train an activity recognition model for detecting these activities using the labeled data and install the activity recognition model onto the loggers that are deployed in the field.

However, since the microcontroller units (MCUs) that can be mounted in small biologgers tend to have limited memory and low computing capability, it is difficult to run computationally expensive machine learning processes on the loggers. In this study, we have developed a computationally efficient animal activity recognition method based on the random forest algorithm that can run on such MCUs. In brief, our method automatically builds a small decision tree for activity recognition that fits in the flash memory of the MCU while maintaining high activity recognition accuracy.

In addition, in order to achieve robust activity recognition, our method also has the following features: (i) robustness to noise, (ii) robustness to sensor positioning, and (iii) robustness to differences in sensor hardware. Robustness to noise refers to the need to handle the varying amount of noise present in sensor data due to differences in how securely the loggers are attached to the animals. Robustness to sensor positioning refers to the need to deal with differences in sensor data collected from different individuals due to variations in the positioning and orientation of the devices. Robustness to differences in sensor hardware refers to the need to handle the differences in sensor data that stem from using data collected from previous years’ hardware when training models for a logger that uses new hardware. We discuss each of these features in the section *Sensor data logger*.

Along with the results reported in this paper, we are also providing open access to some of the software used in this study along with hardware diagrams of the biologgers used, in hopes of assisting other researchers who wish to deploy similar systems in the future. The software includes a labelling tool that can be used to prepare biologger sensor data for use when training machine learning systems and a docker container that includes our algorithm for generating low cost decision trees and scripts for generating the source code needed to run biologgers such as the ones used in this study. This information is available at TBD.

## RESULTS

### Sensor data logger

We begin with a brief introduction to the sensor data loggers used in this study (for more details see Online Methods). Fig. 1 (c) shows a close-up view of the logger, with the camera module located on the far-left end of the logger. Fig. 1 (d) shows an example of the data collected from a chest-mounted logger, with the map displaying the GPS data collected and the two inset images showing frames from foraging activity captured by the device. Fig. 1 (e) shows an example of how these devices were attached in the field. In this example, the logger is attached on the back of the animal, with the camera facing forward and the GPS receiver (white square to the rear of the device) facing the sky. Additionally, in some cases the devices were instead attached to the chest of the birds, in order to improve the camera’s field of view during foraging activities.

Note that because our logger is equipped with a commercially-available MCU and sensors using a simple circuit design (see Online Methods), we believe that reproduction of the logger system using rapid prototyping platforms, such as Arduino, is relatively easy.

### Activity recognition method

#### Overview

Our method is based on supervised machine learning, which can be divided into two main phases: training and testing. This approach assumes that sensor data that corresponds to the data collected by our low-energy sensors can be collected in advance during the training phase. During the training phase, the preexisting sensor data is labelled by biologists to indicate the target activities that should be captured by the loggers’ cameras. This labeled sensor data is then used to train the activity recognition models that will be installed on the biologgers for camera control. The testing phase corresponds to the biologgers’ use in the field, where the model built using the preexisting data is used on board the loggers to recognize target activity in real time using data collected by the loggers’ low energy sensors.

The supervised machine learning method used by our study uses decision trees, in which a hierarchy of simple rules is learned during the training phase that can be used to classify input data vectors during the testing phase based on thresholds learned from the training data. We start with the raw sensor data, which comes from our preexisting dataset in the training phase and from our low-energy sensors during the testing phase. We then divide this raw data into short windows (e.g., 1-second windows), from which we can extract several features from each window that will be used as input for the decision trees. Each window is then represented by a vector of extracted features, with labelled vectors extracted from preexisting data used to train the decision tree during the training phase and unlabeled vectors extracted in real time from low-energy sensors used as input to the tree during the testing phase. In our method, we build these decision trees using a modified version of the random forest algorithm in which we generate trees that minimize the amount of flash memory used for feature extraction on the logging device. The output of the decision tree classifier is then used to control the logger’s video recording, allowing us to conserve the battery power of the logging device by limiting video recording to when we are most likely to capture the target activity.

### Feature extraction

In order to detect an activity of interest, we must first extract features from the raw data collected by our low-energy sensors. In this study, we extract these features from acceleration data and/or GPS coordinates. Fig. 2 (a) and (b) show examples of the GPS and accelerometer data collected by our device.

**Figure 2.**
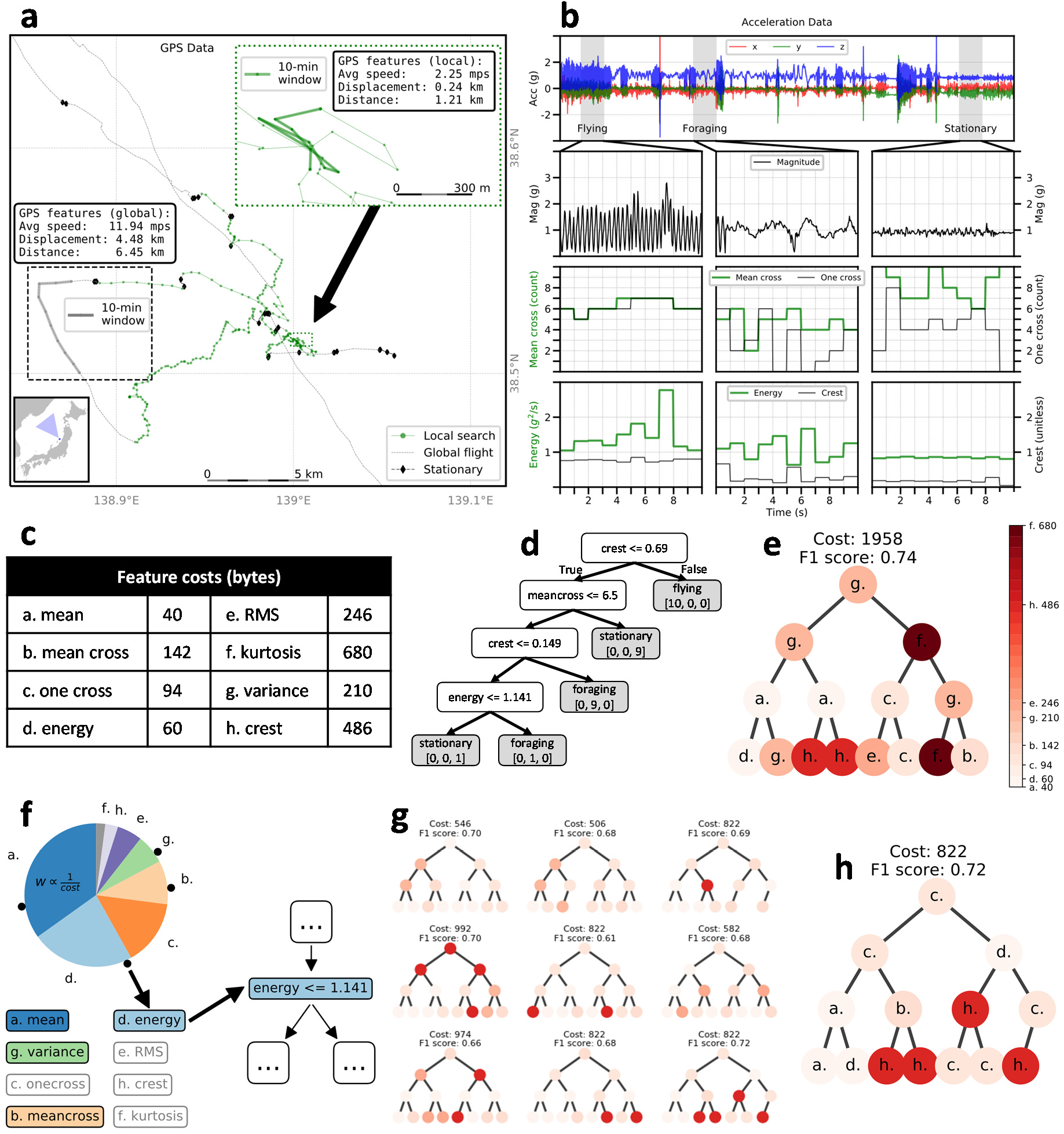
Creating AI models for sensor control onboard biologging devices. (a) Example GPS data collected by our device, collected from a single streaked shearwater from a colony on Awashima Island, Japan. The inset box on the left shows an example of global flight behavior, with example features extracted from a 10-min window of GPS data shown above. (b) Example accelerometer data collected by our device, collected from a single black-tailed gull from a colony near Hachinohe City, Japan. The first row shows raw acceleration data, the second row shows magnitude of acceleration data from 10-sec windows of data corresponding to the behaviors flying, foraging, and stationary, respectively. The bottom two rows show four example features extracted from the magnitude of acceleration data for each 10-sec window. (c) The amount of program memory in bytes used to program each feature extraction function used for the decision tree. (d) Example decision tree generated from the 1-sec segments of feature values shown in the lower two rows of (b). Each white node represents a decision based on a single feature’s value and each grey node represents a final predicted class for the current 1-sec segment of data. (e) Example decision tree generated by a standard decision tree algorithm. (f) Modified weighted sampling of features used in our method. Each feature is randomly selected proportionally to the inverse of their size. (g) Example output from our modified version of the random forest algorithm. Each tree is a candidate low-cost tree for use on a biologging device. (h) Possible final candidate selected from the trees in (g).

The GPS track in Fig. 2 (a) shows the movement of a single bird, with the animal’s positions labeled as belonging to one of three different activity classes: *local search*, *global flight*, and *stationary*. The two inset boxes in Fig. 2 (a) show examples of the global flight and local search activities, along with examples of some features extracted from a 10-minute window of GPS data collected at a rate of one position per minute taken from each example. Comparing the two activities, we can see how such features can capture key differences between the activities. For example, the local search activity is conducted at a lower average speed with low displacement relative to the distance traveled when compared to the global flight activity.

Fig. 2 (b) shows an example of the accelerometer data collected by our device. The data is collected using a sampling rate of 25 Hz, with the net magnitude of acceleration computed for each 3-axis sample and stored in a 25-sample (1-second) buffer in RAM. The conversion from 3-axis data to magnitude values is illustrated in Fig. 2 (b), with the first row corresponding to the raw 3-axis data and the second row corresponding to the converted magnitude data. Features are then extracted from the 1-second windows of magnitude values, with Fig. 2 (c) listing some of the features used in our method. For the full list of features extracted from GPS and accelerometer data see Online Methods.

The acceleration data shown in the first row of Fig. 2 (b) includes three highlighted portions that correspond to the activities: *flying*, *foraging*, and *stationary*. The third row of Fig. 2 (b) shows the magnitude data for each activity, while the third and fourth rows show examples of the features that are extracted from 1-second windows of magnitude data in our method. Note that each horizontal segment in the stepped lines in the third and fourth rows correspond to the single value extracted for the 1-second window covered by the horizontal segment. Comparing the three activities, we can again see how these features capture key characteristics of each activity allowing us to distinguish between the activities based on a few key values derived from each window of data.

### Classification

Using the features extracted from the GPS or accelerometer data, we then construct a decision tree that can be used to classify each segment of data into an activity class. Fig. 2 (d) shows an example of such a tree that was constructed using Scikit-learn’s decision tree algorithm (Pedregosa et al. 2011) using the magnitude-based features shown in rows three and four of Fig. 2 (b). The white nodes in this tree show the rules used to classify each instance of data based on the features extracted from the sensor data, while the grey leaf nodes show the classes assigned based on those rules. Each leaf node also lists the support for each class at that node, with the three values listed (e.g., [10, 0, 0]) corresponding to the number of instances of training data classified at that node from the classes flying, foraging, and stationary, respectively. In this example the support values from all the leaf nodes sum to 30, which correspond to the 30 segments of training data taken from rows three and four of Fig. 2 (b).

When classifying a new 1-second window of data, we simply start at the root node of the tree and compute each feature encountered until we reach a leaf node that assigns the most likely class for the data. For example, consider the case where we need to classify one of the 10 1-second windows from the *Flying* portion of Fig 2 (b), i.e., the data shown in the left-most chart of each of rows two through four. Starting at the root node of the example decision tree in Fig. 2 (d), we see that the first rule used during classification checks the *crest* feature using a threshold of 0.69. Given that the *crest* values for our *Flying* data are all greater than 0.69, we would follow the *False* path from that node, immediately reaching a leaf node that assigns the *Flying* class to the data segment.

Looking at this example tree, we can also better understand two potential benefits of decision trees when used with MCUs. The first is their small size, which is due to how their logic can be implemented as a series of nested if-else statements. This allows their models to be stored using only a minimal amount of flash memory (as opposed to models generated by other techniques such as SVM which typically consume too much space for use on MCUs). The second is their potential for minimizing the energy used by the device during recognition. This comes from how each input data segment only follows a single path through the tree, meaning that the MCU needs only to extract features as they are encountered in the path taken through the tree, minimizing the feature extraction processes run for each input vector.

### Feature costs

Standard decision tree algorithms, e.g., Scikit-learn’s default algorithm, build decision trees that maximize classification accuracy with no option to weight the features used in the tree based on a secondary factor such as memory usage. This can be an issue when running the classifier on an embedded device, where the total amount of flash memory available can be limited (e.g., 32 kB). Fig. 2 (e) shows an example of a decision tree built using Scikit-learn’s default algorithm using a full dataset of acceleration data, which results in a total memory footprint of approximately 1958 bytes. While this tree technically fits into our biologger’s limited flash memory, its large size reduces the memory available for other system functions needed to operate the logger’s sensors and write sensor data to long-term storage.

The memory size of the tree in Fig. 2 (e) was estimated based on the feature sizes listed in Fig. 2 (c), with the letters used to label each node in the tree indicating which feature from Fig. 2 (c) was used at that node. While freely choosing a combination of several of these features results in an accurate decision tree, it may also be possible to achieve good results when using only a subset of these features. By restricting the use of the costliest features, e.g., kurtosis, it may be possible to reduce the size of the resulting tree while achieving similar accuracy.

### Reduced cost decision tree

Given the need to minimize the size of the feature extraction functions used by decision trees when run on devices such as biologgers, this study proposes a method for automatically generating low-cost decision trees that is based on the *random forest* algorithm (Breiman, L. 2001). The random forest algorithm is a decision tree algorithm that generates multiple unique decision trees from a single dataset by restricting the features made available when creating each node in a tree to a randomly selected subset of the features. In the original random forest algorithm, several trees are generated in this way and are then combined for use as an ensemble classifier. Our method modifies the original algorithm by using weighted random selection of the features for each node, with each feature extraction function assigned a weight proportional to the inverse of its size. The resulting algorithm generates randomized trees that are less likely to incorporate features that require more space in flash memory while still attempting to maximize classification accuracy using the remaining features. We then select a single tree from among the several randomized trees generated for use on our device.

Fig. 2 (f) shows the process used when generating nodes in a decision tree using our method. We start by assigning each feature a weight that is proportional to the inverse of its weight. For example, *mean* uses only 40 bytes of flash memory and so is assigned a relatively high weight of 0.35, while *kurtosis* uses 680 bytes of flash memory and so is assigned a weight of 0.02. We then use these weights to perform weighted random selection (without replacement) of the features to select which features to use when creating a new node in the tree. In this example, we have randomly placed four dots along the perimeter of the pie chart, signifying a random selection of the features *mean*, *variance*, *mean-cross*, and *energy*. We then select the best candidate feature from amongst these randomly selected features, which in this example is energy. This feature is then used to create the next node in our decision tree, shown as the node “energy <= 1.141” on the right side of Fig. 2 (f).

Using our method for weighted random selection of nodes described above, we are then able to generate randomized trees that tend to use less costly features. When generating these trees, we can easily estimate the size of each tree generated based on the sum of sizes of all features used in the tree and can set a threshold size for which all trees above the threshold are discarded. Fig. 2 (g) shows an example batch of trees output by our method where we have set a threshold size of 1000 bytes. These trees were generated from the same training and validation data as was used for Fig. 2 (e). We can then select a single tree from among these trees that gives our desired balance of cost to accuracy. In this example, we have selected the tree illustrated in Fig. 2 (h) based on it having the highest accuracy among this batch of trees. Comparing Fig. 2 (h) to (e), we can see that our method was able to generate a tree that is 42 percent the size of (e) while maintaining close to the same accuracy.

### Other functionalities of our logger

Our method also incorporates several functionalities that enable robust activity recognition in the conditions encountered during this study. First, we address the need for noise robustness, due to the varying amount of noise that can be introduced into the sensor data stemming from how the loggers must be loosely attached to the birds via taping the logger to the birds’ feathers. We achieve this through data augmentation during the training phase, in which we train our models on multiple versions of our training data that each are altered by adding varying levels of random artificial noise. Next we address the need for robustness to sensor positioning, which stems from how loggers can be attached to birds at different positions and orientations, such as some loggers having been placed on the birds’ backs to maximize GPS reception while others were placed on the birds’ chests to improve the camera’s view of the animals’ feeding. We achieve this by converting all 3-axis accelerometer data to net magnitude of acceleration values, removing the orientation information from the data before use in our activity recognition models. Finally, we address the need for robustness to differences in sensor hardware that stems from how the biologgers used in this study must be trained using accelerometer data collected from hardware used in previous years’ research. We achieve this by running an online conversion of the sensor data collected on our biologger to downsample our sensor’s 16-bit resolution data to match the 8-bit resolution data collected in previous years prior to using the data in our activity recognition models. Further information about these functionalities can be found in the Online Methods.

### Performance of Proposed Method

We evaluated the proposed method by using it to control the video recorded by the biologgers described in the section *Sensor data logger* when attached to black-tailed gulls from a breeding colony located on Kabushima Island at Hachinohe, Japan. Along with the proposed method, we also deployed one logger using a naive method, in which the logger was programmed to activate the camera in 15-minute intervals. All loggers (naive and proposed) ran the camera for a set 1-minute window after each activation. Altogether 11 loggers were used, with 10 loggers running the proposed method and 1 logger running the naive method. A total of 212 1-minute videos were collected by the loggers, with 185 videos collected using the proposed method and 27 videos collected using the naive sampling method.

The proposed method was trained to activate the cameras during possible foraging activity, which we identified based on abnormal movements during flight that seemed to correspond to diving behavior. These abnormal movements were detected by extracting features from 1-second windows of acceleration data. Additionally, camera activation was limited to movements detected during flight activity by only activating the camera when the bird’s movement had recently been classified as flying prior to being classified as foraging, i.e., flying had been detected within the previous five seconds. The acceleration data used to train the decision trees used in our method was collected in the previous year from birds at the same colony using Axy-trek logging devices^1^.

Fig. 3 gives an overview of the results for the black-tailed gulls. Fig 3 (a) and (b) show GPS tracks that give an overview of the video data collected by the proposed method and the naive method, respectively. The portions of the tracks highlighted in green show where video data was collected on possible or confirmed foraging activity, while the sections highlighted in grey show where video was collected on non-foraging activity. While only one logger was run using the naive sampling strategy, its results highlight the issue with such a method, with the logger quickly depleting its battery recording videos on and around the nesting area, greatly reducing the range of collection when compared to the devices using event-based camera activation.

**Figure 3.**
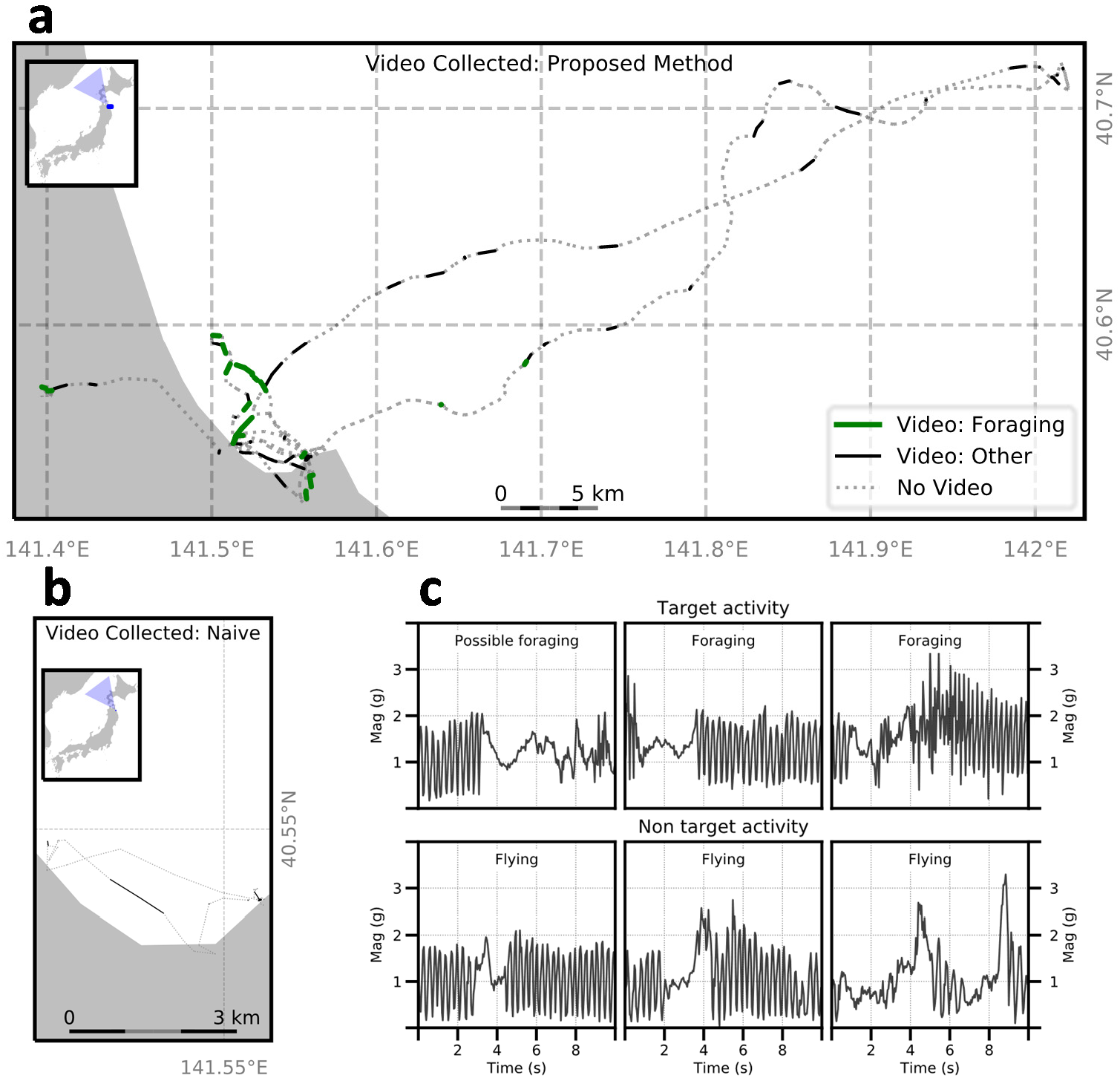
Results of AI video control for black-tailed gull. (a) GPS tracks collected by biologgers using the proposed method. Green highlighted sections represent successful video collection of foraging behavior. Grey sections represent video collection of non-target behavior. (b) GPS tracks collected by biologger using the naive method. (c) Examples of acceleration data (shown as magnitude of acceleration) collected around the time of video camera activation on biologgers using the proposed method. Top row corresponds to videos containing target behavior while bottom row corresponds to videos with non-target behavior.

Fig. 3 (c) shows six examples of the acceleration data that was collected surrounding the time of camera activation by the proposed method, with each chart showing 10 seconds of net magnitude of acceleration data corresponding to a single example. The first row shows three examples of foraging and possible foraging activity, in which the camera was correctly activated based on the birds’ movements, while the three examples in the second row show non-target (flying) activity in which the camera was incorrectly triggered. Note that the camera is activated based on a 1-second window of data, which corresponds to a window extracted from the area around the 2 to 4 second mark for each example. The exact timing is not known due to a short delay (about 3 to 4 seconds) between when the camera is triggered and when it starts recording data. As is shown in these charts, while acceleration data can be used to detect the target activity, it is difficult to avoid false positives due to the similarity between the target activity and other anomalous movements in the sensor data. Furthermore, due to the camera delay, it is not possible to film short actions that do not last longer than the camera delay or repeat within the 1-minute recording window. Some of the camera activations determined to be false positives in these results may have been such one-time actions that were not captured due to the delay.

The 212 videos collected by the biologgers were evaluated by the biologists participating in this study, with each video classified as belonging to the classes: foraging, possible foraging, flying, and stationary. Of the 27 videos collected by the naive method, none contained any target activity, with 3 videos containing flying activity and 24 videos containing stationary activity. In contrast, of the 185 videos collected by the proposed method, 58 contained target activity (5 confirmed foraging and 53 possible foraging) and 127 contained non-target activity (86 flying and 41 stationary), giving the proposed method a precision of about 0.31. Of particular interest were five target activity videos which captured images of the black-tailed gulls feeding on insects, both over land and over the sea. (**Supplementary Videos 1 & 2**)

Along with the evaluation done by the biologists, we also analyzed the performance of the biologger by first fully labelling the low-energy sensor data (i.e., accelerometer data) collected by the biologgers and then computing the precision, recall, and f-measure for the 1-minute windows of sensor data that corresponded to the 212 videos collected by the logger. Based on this full labelling of the data, we computed the estimated distribution of the activities in the sensor data and found that the target activity (foraging) comprised only about 2 percent of the 6,616 total minutes of data collected, with 10 percent corresponding to flying activity and the remaining 88 percent corresponding to stationary. The proposed method achieved a precision of 0.27, a recall of 0.56, and an f-measure of 0.37 based on this full labelling. The naive method was again determined to have not collected any target activity, and so received a 0 for all three scores. Meanwhile, the proposed method was able to capture about half of the estimated windows of target activity (recall 0.56) and achieved a precision of 0.27, which is well above the expected precision of 0.02 for a naive sampling method when the target comprises only 2 percent of the dataset.

## DISCUSSION

Several previous studies involving biologgers have introduced trigger mechanisms that can be used to control when high-cost sensors are activated, with many of these studies focusing on controlling animal-borne cameras such as the one used in this study. (Troscianko et al. 2015) introduced a programmable animal-borne camera that incorporated an internal clock, allowing their camera to only be activated at set times of day. Both (Beringer et al. 2005) and (Goldbogen et al. 2017) incorporated light sensors to prevent their cameras from being triggered during periods of darkness. (Boness et al. 2006) ensured that their camera only recorded the animal when it was at sea using a saltwater switch. (Watanuki et al. 2007) and (Volpov et al. 2015) took this a step farther by incorporating a depth sensor, allowing their cameras to only trigger when an animal surpassed a predefined depth threshold. (Nishiumi et al. 2018) deployed devices with two acceleration sensors, using a low-cost (low-frequency) acceleration sensor to activate a second high-cost (high-frequency) acceleration sensor when a preset threshold had been surpassed. Finally, (Brown et al. 2012) measured the variance from their low-cost acceleration sensor to dynamically adjust the sampling rate of their high-cost GPS sensor based on predetermined threshold values. In each of these previous studies, the readings from the low-cost sensors were only compared to preset thresholds when determining whether to activate a high-cost sensor. Such methods are only suitable for coarse-level characterizations of behavior such as differentiating between underwater activity versus surface activity. In contrast, our proposed method can be used to distinguish between complex behaviors at a finer scale, allowing biologists to target a specific target behavior.

This is the first study to our knowledge to deploy AI in animal-borne data loggers. Wild animals represent one of the most extreme environments in which AI works in terms of limited space and harsh conditions. We anticipate our work will provide motivation for more widespread adoption of AI techniques on biologgers, both for intelligent sensor control and intelligent onboard data processing. Such techniques can not only be used to control what is collected by such devices, but also what is transmitted off the devices, such as is done by satellite relay tags (Cox et al. 2018). The combination of IoA (Internet of Animals) and AIoA (AI on animals) would enable biologists to answer a number of scientific questions about wild animals and obtain important information for their conservation.

## DATA COLLECTION

The research at Awashima Island was conducted with permits from the Ministry of the Environment, Japan. The protocols at Kabushima Island were approved by the Agency for Cultural Affairs, Japan and the Aomori Prefectural Government. All field protocols were approved by the Animal Experimental Committee of Nagoya University.

## ACKNOWLEDGEMENTS

We thank Rory P. Wilson, Yasue Kishino, and Kazuya Murao for suggestions and comments on this work. This work was supported by JSPS Kakenhi JP16H06536, JP16H06539, and JP16H06541.

## AUTHOR CONTRIBUTIONS

J.M.K. performed method design, software implementation, data collection, data analysis, and manuscript writing. H.S., S.M., and Y.M. performed data collection and data analysis. M.S. performed software implementation and data collection. T.M. conceived and directed the study, and performed method design, data collection, data analysis, and manuscript writing. J.N. performed hardware design. K.Y. performed data collection, data analysis, and manuscript writing.

## Footnotes

1 http://www.technosmart.eu/axytrek.php

